# Rapid loss of fine motor skills after low dose space radiation exposure

**DOI:** 10.1101/2022.04.01.486651

**Authors:** Ashley A Blackwell, Arriyam Fesshaye, Alyssa Tidmore, Rami I Lake, Douglas G Wallace, Richard A Britten

## Abstract

Sensorimotor function, motivation, and attentional processes are fundamental aspects of behavioral organization during skilled tasks. NASA’s planned expedition to Mars will expose astronauts to space radiation (SR) that has the potential to impair performance in mission critical tasks. Impairments in task accuracy and movement kinematics have been previously reported during string-pulling behavior ~7 months after SR exposure. If similar SR-induced sensorimotor deficits emerge at earlier times, then astronauts may have compromised in-flight performance disruptions while performing skilled tasks in critical situations, such as when manipulating controls or performing seat egress. Due to the possibility that such performance losses may compromise mission success, it is critical to determine if sensorimotor, motivation, or attentional deficits occur acutely after SR exposure at a time point that corresponds to in-flight performance. Male Wistar rats were thus exposed to either 10 cGy simplified galactic cosmic radiation (GCRsim), 10 cGy ^4^Helium (^4^He), or no radiation at all (Sham), and string-pulling behavior was assessed approximately 72 hours later. Following exposure to SR, rats (^4^He) took more time to approach the string to initiate string-pulling behavior and to pull in the string to reach the Cheerio (^4^He and GCRsim) relative to Sham rats. ^4^He-exposed rats also exhibited a greater number of misses and less contacts relative to both Sham and GCRsim-exposed rats. Further, rats exposed to ^4^He demonstrated less concentrated reach endpoints with both the left and right hands compared to GCR-exposed rats. This work suggests that sensorimotor function and motivation and/or attentional processes were impaired 72 hours after ^4^He-radiation exposure.

## 1. Introduction

Astronauts will soon travel further into space than ever before exposing themselves to unavoidable deep space radiation (SR). The radiation spectrum found in deep space, frequently termed Galactic Cosmic Radiation (GCR), is composed of highly energetic, high mass (Z ≤ 28) charged ions. Current estimates suggest that astronauts will be exposed to ~30-40 centigray (cGy) of SR on a three-year mission to Mars and back [1]. While SR exposure contributes to central nervous system deficits that impair several aspects of mission critical performance, including multiple cognitive domains [for review see 2, 3–4], the impact of SR on sensorimotor function is not as defined. The few studies characterizing SR effects on sensorimotor function to date have primarily focused on gross aspects of performance [5–8]. This previous work also used SR ions that contribute to less than 1% of the dose within the Local-Field spectrum: the SR spectrum that the internal organs of astronauts will receive within the spacecraft. The majority of the physical and dose equivalent SR dose will arise from high energy, low mass (Z < 15) particles [9–10].

Recent advancements in the generation of simulated SR ion beams have allowed for the development of more sophisticated exposure models. From a practical perspective, it was only possible to deliver one ion at a time during irradiation exposure but advancing technology has provided the opportunity for multiple ions to be delivered during one exposure session (i.e., 6-ion GCRsim). The relative dose contribution of the 6-ion GCRSim beam is 74% protons, 18% ^4^He, 6% ^16^Oxygen, 1% ^28^Silicon (Si), and 1% ^56^Iron [11]. Only a handful of studies have been conducted to evaluate the effects of GCRsim on cognition [12–16], and none of this work has characterized sensorimotor function.

The string-pulling task provides a robust assessment of sensorimotor function, including fine motor control. During this behavior, both rodents and humans engage in highly organized bimanual hand-over-hand movements to pull in a string [17–18]. There has been a single study to establish the impact of SR exposure alone on fine motor skills [19]. This work showed that exposure to 5 cGy ^28^Si significantly disrupted string-pulling behavior ~7 months after irradiation in rats that had no obvious loss of executive function performance. While the loss of fine motor skills later in life would be an issue for astronauts upon their return to Earth, in-flight disruptions can have serious consequences for mission success. When these fine motor skill deficits initially emerge following SR exposure is currently unknown. Therefore, the current study established if SR-induced sensorimotor performance deficits were present at a time point consistent with astronauts on-board a space craft. To do so, a side-by-side comparison of string-pulling behavior was conducted for Sham rats relative to rats exposed to 10 cGy of either GCRsim that mimics the LET characteristics of the Local Field spectrum or 250 MeV/n ^4^He particles at an acute time point (72 hours).

### 2. Methods

### 2.1. Subjects

Male Wistar proven breeder rats (HlaR(WI)CVFR; Hilltop animals, Inc., Scottsdale, PA, USA) were used in the current study. Following one week of acclimation, rats were weighed and implanted with ID-1000us RFID transponders (Trovan Ltd, United States) for easy identification throughout the study. Throughout the study, all rats underwent a treadmill exercise regimen for 25 minutes at a rate of 15 cm/s twice per week, except while at Brookhaven National Laboratory (BNL). All rats were prescreened for cognitive performance in an attentional set shifting task (ATSET) as previously described [20]. Rats that passed the ATSET prescreening were pair-housed and shipped to BNL for SR exposure and string-pull testing. Upon arrival at BNL, pair-housed rats were left undisturbed to acclimate to the environment in consistent vivarium temperatures (20 to 21 °C) on a reverse 12-hour light/dark cycle and provided food and water *ad libidum*. The Institutional Animal Care and Use Committee at BNL and Eastern Virginia Medical School approved all procedures described in this experiment and all guidelines set by the Office of Laboratory Animal Welfare were followed.

### 2.2. Procedures

After one week of acclimation at BNL, rats that continued to be pair housed began food restriction with *ad libitum* access to water. Then, rats were habituated for the string-pulling task which involved single housing rats in a standard Plexiglas home cage (46 cm x 26 cm x 26 cm) for ~2 h with exposure to 20 strings (0.2 cm diameter, 100% cotton, unscented) of varying lengths (30 cm to 150 cm), half with a Cheerio tied at the end, since partial reinforcement yields the highest response rate, as described previously [21]. Only rats that successfully pulled in strings during habituation were included in the string-pulling test (N = 59). Following habituation, rats were randomly a363ssigned to three test groups: Sham (n = 11), 10 cGy ^4^He (n= 24), or 10 cGy GCRsim (n = 22). Radiation exposure occurred at NASA Space Radiation Laboratory (NSRL) when the rats were ~7 months of age and involved placing rats individually into small red irradiation jigs (see Figure 1A) as to continue the rats’ dark cycle throughout irradiation exposure. Rats were irradiated with 10 cGy of either 250 MeV/n ^4^He at a dose rate of <1 cGy/min totaling ~12 min exposure period or 6-ion GCRsim at a dose rate of 0.5 cGy/min totaling ~20 min. Sham rats were placed in the irradiation jigs for the same duration as the SR-exposed rats, without any exposure to irradiation.

**Figure 1:**
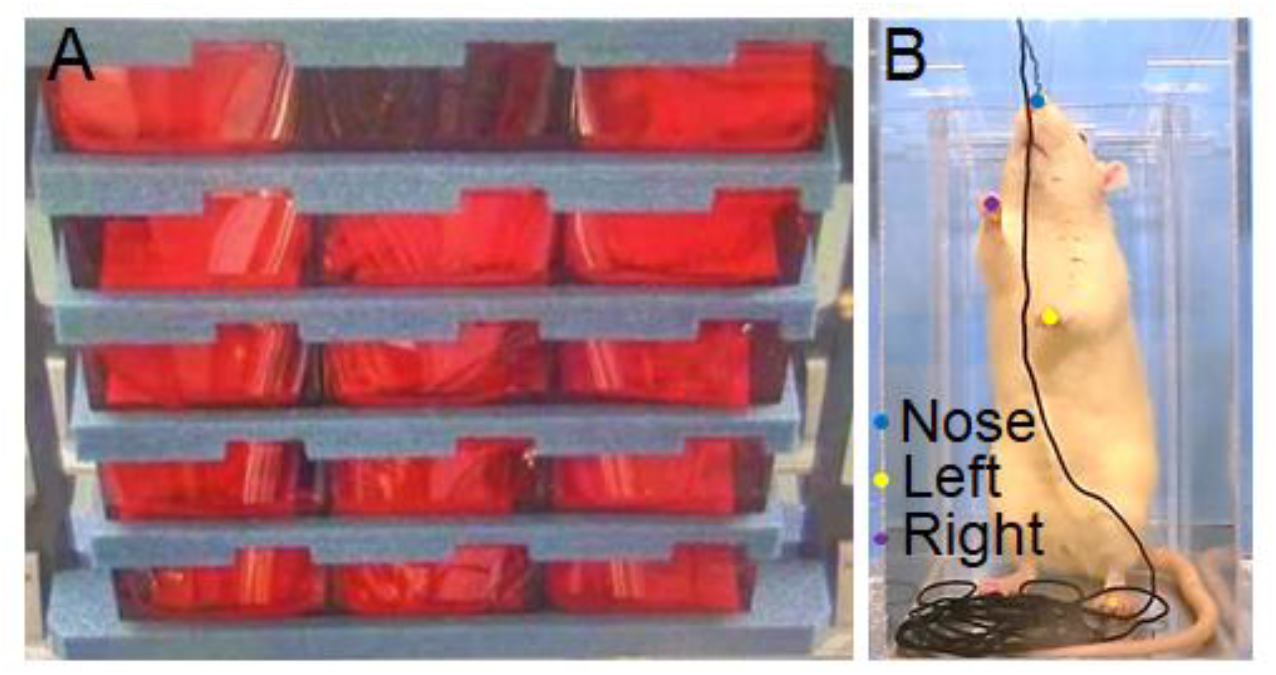
Rats were placed in small red jigs during irradiation and Sham exposure conditions (A). A rat is shown inside the string-pulling apparatus with the string draped back to provide a clear view of the rat (B). The rats body parts were labeled by DLC (V2.1 [22]; nose: blue, left: yellow, and right: purple).

Approximately 72 hours after radiation exposure, rats were tested in the string-pulling task. The string-pulling apparatus consisted of a clear rectangular Plexiglas box (28 cm X 14 cm X 26 cm) with a Plexiglas insert that required rats to stay toward the front of the apparatus and a lid that was placed on a table approximately 1.5 m above the floor in a small room. A string-pulling session consisted of 6 trials to pull in a 59 cm long string baited with a half piece of Cheerio tied at the end. The string was hung in front of the apparatus and draped back to provide a clear view of the rat while pulling in the string (see Figure 1B). Rats remained in the apparatus for all 6 trials, and the apparatus was thoroughly cleaned between rats. A session ended if the rat failed to engage in string-pulling behavior for 20-minutes, and then, the rat was tested the next day. A Canon HD video camera (model #: XA30) set at 1:1000 shutter speed and 59.98 frames per second was positioned perpendicular to the wall of the apparatus to record string-pulling behavior for offline analysis.

### 2.3. General performance measures

The amount of time it took rats to approach and to pull in strings during habituation was recorded. This pre-exposure setup did not allow for the characterization of misses and contacts or kinematics, since rats were habituated in standard Plexiglas home cages (46 cm x 26 cm x 26 cm). Therefore, this analysis provided a basis for pre-versus post-exposure comparisons of approach time and pull duration only.

During testing in the string-pull task, once placed in the apparatus, rats first detect the string and then engage in string-pulling behavior to retrieve a piece of Cheerio tied at the end. The amount of time it took rats to first engage in string-pulling behavior after the string was draped in the front of the apparatus was recorded as the approach time. Pull duration is a measurement of the amount of time it took rats to fully pull in the string once string-pulling behavior was initiated until the Cheerio tied at the end was reached. Approach time and pull duration were evaluated as general measures of performance that may be influenced by motivation [21] and/or attentional processes.

### 2.4. Characterization of movement accuracy

Rats engage in grasps to contact the string and occasionally miss, or fail, to contact the string when attempting to grasp it with both hands and the mouth. Total attempts (i.e., contacts and misses) with the left-and right-hands and the mouth were summed across string-pulling bouts, and the proportion of misses and contacts relative to total attempts, termed movement accuracy ratio, were evaluated between groups. Values closer to one represent a greater number of misses while values closer to zero denote more contacts. Misses provide one assessment of fine motor control that may occur due to either issues with hand dexterity, such as an inability to grasp the string, or to errors in distance and direction estimation related to kinematics. Kinematic components

DeepLabCut (DLC 2.1) [22] was used to track both the hands and nose during bouts of string-pulling behavior (see Figures 2 and 3 for labeled DLC screenshot). A network was trained in DLC using one video from each rat; twenty random frames from each video were extracted via k-means extraction method and manually labeled. Once the network was trained, two other videos for each per rat were labeled for a total of three videos per rat for kinematic analyses. After all videos were labeled with DLC, the XY output was screened for errors, including missing or incorrect labels. DLC provides likelihood values, ranging from zero to one with zero being less confident and one being most confident, as a prediction of accuracy for the labeled body parts. Likelihood values less than 0.050 were generated only when a body part was missing a label but not when an error occurred in labeling. Two methods were used to deal with these different types of errors. First, the XY data corresponding to either type of error was deleted and averaged using the accurately labeled data points before and after the errors. Second, if too many DLC labeling errors occurred in a row to produce accurate movement, then the string-pulling bout was broken up into two separate parts and averaged across each part to include all data.

**Figure 2:**
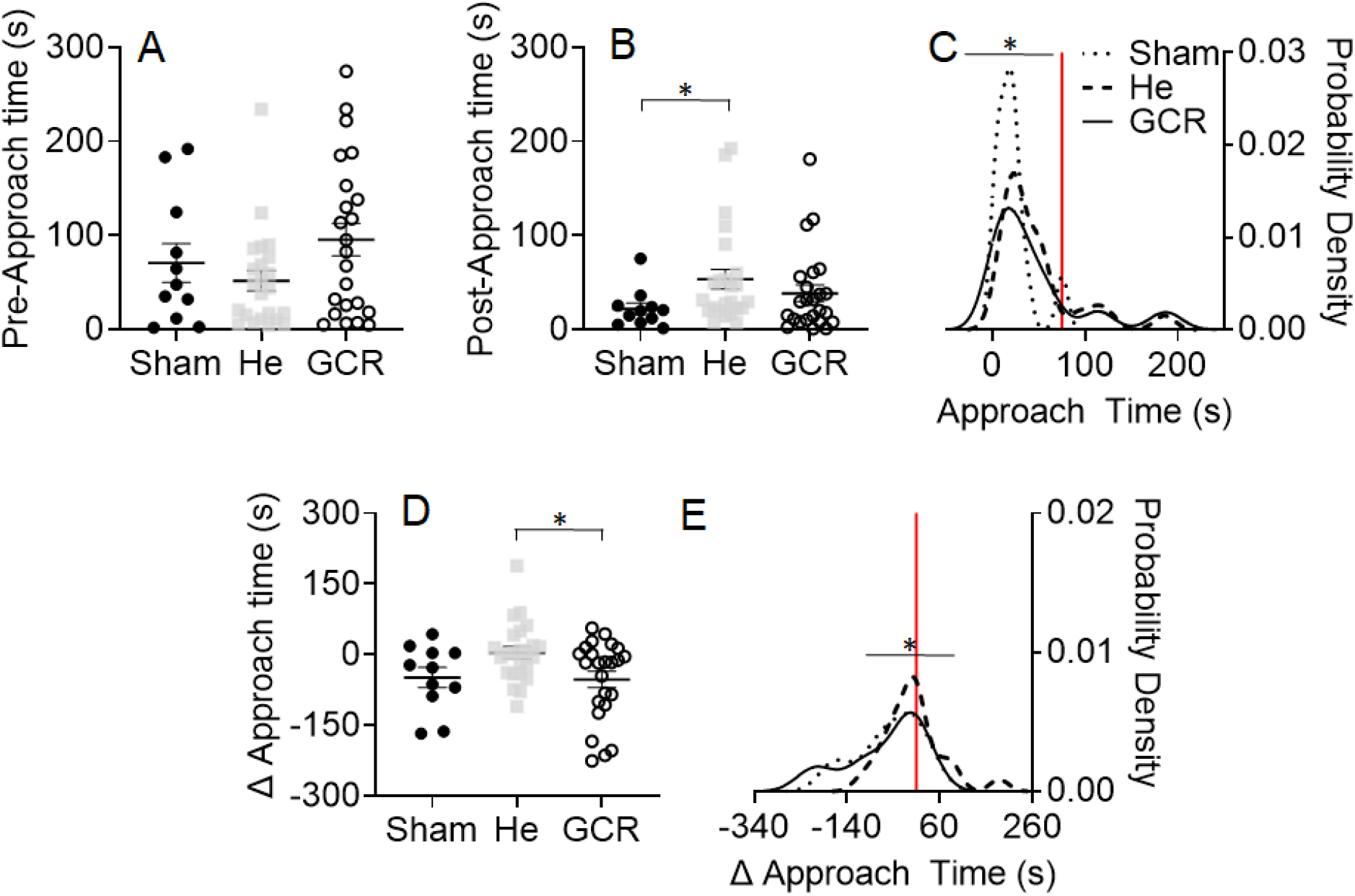
The time it took rats to approach time string to begin pulling is shown before irradiation (A). After SR exposure (B) significant differences were observed between ^4^He-exposed and Sham rats. The probability density profile generated from the KDE analysis of post-SR approach time revealed further significant group differences for each SR-exposed group relative to Sham rats (C). The change in approach time across conditions depicted by group significantly differed between irradiated groups with 4He rats taking longer to approach the string relative to GCRsim rats (D). Further, probability density profiles generated from the KDE analysis of the change in approach time yielded significant group differences for each SR-exposed group relative to Sham rats and between irradiated groups (E). *p < 0.050

**Figure 3:**
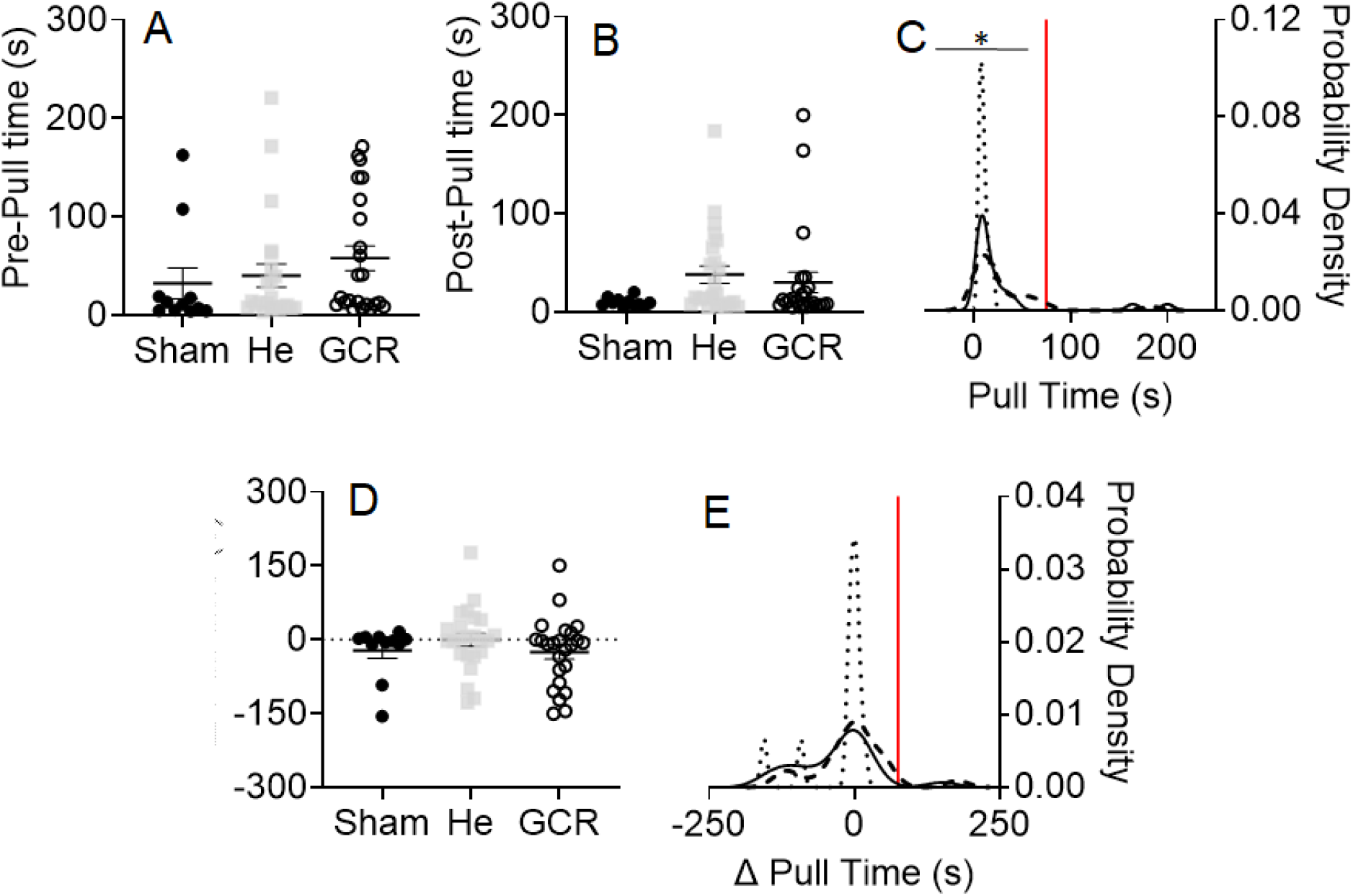
The amount of time it took SR rats to pull in the string to reach the Cheerio tied at the end was similar between groups before (A) and after (B) irradiation exposure. However, when the probability density profile was generated using the KDE analysis, significant group differences were observed between each SR-exposed group compared to Sham rats (C). Assessments of the change in pull duration across conditions failed to reveal significant group differences for traditional analyses (D) and for KDE generated probability density profiles (E). *p < 0.050

The screened XY coordinates generated from tracking with DLC were used to evaluate movement organization of both hands and the nose during the string-pulling task. Bouts of string-pulling behavior was segmented into reaches and withdraws to evaluate general and specific kinematic performance measures of movement organization. Distance traveled (cm) and peak speed (cm/s) were characterized as general kinematic measures. Path circuity was a specific measure used to characterize how direct the hands traveled during reach and withdraw trajectories. Values generated closer to zero are indicative of more circuitous paths, and values approaching one represent more direct paths.

Circular statistics was used to evaluate the concentration (first order) of reach and withdraw endpoints and their average heading directions (second order) [23]. Concentration is defined as the strength of reach and withdraw endpoint clustering, with values closer to zero representing randomly scattered endpoints, and values closer to one indicating tightly clustered endpoints. Heading direction refers to the average directional heading trajectory of reach and withdraw endpoints and ranges from zero to 360 degrees. Reaches are typically directed upward toward 90 degrees to grasp the string at or just after the peak of the reach, while withdraws are oriented downward toward the midline of the body near 270 degrees to pull the string down into the apparatus [24].

Rats engage in mouth contacts to pull in the string and use their nose to track the string while pulling it into the apparatus. Therefore, nose kinematics were characterized across the entire string-pulling bout. The peak speed (cm/s) that the nose traveled was compared between groups along with the range of nose movement (cm) in both the x- and y-axes. These kinematic measures were used to investigate general movement organization of the nose during bouts of string-pulling behavior.

### 2.6. Statistical Analyses

Data was visualized graphically with QQ and frequency plots, and Shapiro-Wilk and Levene’s tests (p < 0.050) were used to check normality and homogeneity, respectively, for each measure. Normality was violated for general measures of performance, including approach time and pull duration; thus, non-parametric statistical analyses were used to compare groups. Mann Whitney tests were conducted with the 95% confidence intervals reported to compare each SR group to the Sham rats, since this test does not require normal distribution of data and equal variances.

Kernel Density Estimation (KDE) was calculated using the Kernel Density Estimation program in the Free Statistics Software (v1.2.1) [25] with the Gaussian distribution as Kernel; the bandwidth set to zero to allow the algorithm to select the optimal bandwidth to fit the data based on the inter-quartile range. Since KDE produces values outside of the domain range (i.e., negative values generated for approach and pull measures), when the string-pulling data was modeled, the KDE generated probability distribution profile was trimmed of data below zero seconds for all three groups. The resulting probability profile function was used to establish the 5^th^ worst percentile performance metric of the sham rats which provided a basis of comparison to determine how SR exposure altered the area under the curve that exceeded that level of performance. Fisher’s exact test (p < 0.050) was used to assess group differences between each SR group and Sham rats. An advantage to KDE analysis is that the data generated by the probability distribution profile may be used to calculate a risk ratio and numbers needed to harm which are of most importance to NASA and cannot be conducted with raw data. The risk ratio is defined as an increase or decrease in absolute risk level, while numbers needed to harm refers to the number of individuals that must be exposed to SR for one of them to present an adverse reaction from exposure [Elliott et al., 2020; Suchmacher, Geller, 2021]. Therefore, this data was also entered into a risk ratio calculator to determine the level of risk and the numbers needed to harm [26]. GraphPad Prism 9 statistical software was used for Mann Whitney and Fisher’s exact non-parametric tests.

Assumptions were not violated for any other performance measures. Therefore, an ANOVA was used to investigate the main effects of group on movement accuracy ratio. For reach and withdraw kinematic subcomponent analyses, repeated measures ANOVAs were used to evaluate group differences between the left and right hands. Alpha was set to p < 0.050 and partial eta squared (η^2^_p_) was reported as a measure of effect size. Tukey HSD (p < 0.050) was used to further investigate group differences for post-hoc analyses. GraphPad Prism 9 was also used for parametric statistics with confirmatory analyses conducted using Jasp 0.14.0 statistical software (University of Amsterdam).

## 3. Results

### 3.1. General performance measures

#### 3.1.1. Approach time

All rats engaged in string-pulling behavior during habituation before irradiation (i.e., 18 out of 20 strings pulled in), but two rats failed to approach the string and engage in string-pulling 72 hours after exposure to GCRsim. For the rats that performed in the string-pulling task after SR exposure (N = 57), ion-specific effects were observed on the amount of time it took to approach the string to initiate string-pulling behavior (see Figure 2A). Approach time was investigated prior to radiation exposure, 72 hours following irradiation, and the chfange between the two sessions to determine changes in performance.

While no significant Group differences were observed with an ANOVA conducted on pre-irradiation approach time [F (2, 57) = 2.302, p = 0.110, η^2^_p_ = 0.079] between Sham (M = 70.578, SD = 68.229), ^4^He (M = 51.685, SD = 52.392), and GCRsim groups (M = 95.403, SD = 83.045), rats exposed to ^4^He (M = 53.413, SD = 51.516) exhibited a significantly longer approach time compared to Sham (M = 21.980, SD = 20.368) rats [Mann Whitney: p = 0.020, CI: 2.580 to 38.970] after SR exposure. However, GCRsim (M = 40.056, SD = 43.683) rats failed to differ from Sham rats [Mann Whitney: p = 0.221, CI: m11.830 to 44.700], and irradiated rat failed to differ from each other [Mann Whitney: p = 0.184, CI: 26.630 to 4.050]. Exposure to ^4^He specifically increased approach time relative to Sham rats 72 hours later (see Figure 2B).

Next, a KDE analysis of approach time calculated the 5^th^ worst percentile for Sham rats to be 75.20 s (see Figure 2C). The percentage of SR-exposed rats that had performance below the 5^th^ percentile of the Sham rats’ performance was 18.4% for ^4^He- and 22.2% for GCRsim exposed rats. Fisher’s exact tests revealed significant differences for ^4^He-[p = 0.0067] and GCRsim [p = 0.0007] rats relative to Sham rats. Both SR groups had a significant percentage of rats whose performance was below the 5^th^ worse percentile of Sham rats but failed to differ from one another [Fisher’s exact test: p = 0.596]. Then, the KDE generated data that was entered into a risk ratio calculator determined both level of absolute risk increase and numbers needed to harm for ^4^He rats to be 13.59% (CI: 4.81% to 22.36%) and 7 (CI: 4.5 to 20.8) and GCRsim rats to be 17% (CI: 7.83% to 26.17%) and 6 (CI: 3.8 to 12.8), respectively. Approximately every 7 rats exposed to either ^4^He or GCR will result in one rat taking longer amounts of time to approach the string.

The difference in approach time post-relative to pre-SR exposure was investigated as a function of Group (see Figure 2D). An ANOVA conducted on this change in approach time revealed a significant effect of Group [F (2, 56) = 4.148, p = 0.021, η^2^_p_ = 0.133]. ^4^He (M = 1.878, SD = 63.586) rats took a longer amount of time to approach the string than GCRsim (M = −57.756, SD = 83.376) rats post-compared to pre-exposure, while Sham (M = −48.597, SD = 69.995) rats failed to differ from either irradiated group. In addition, KDE analyses calculated the 5^th^ worst percentile of Sham rats’ performance to be 9.13 seconds for the change in approach time from post-to pre-exposure conditions with 42% of ^4^He and 25% of GCR rats failing below this 5^th^ percentile (see Figure 2E). Fisher’s exact tests revealed significant differences for ^4^He- [p < 0.0001] and GCRsim [p = 0.0007] rats relative to Sham rats. Further analysis between irradiated groups also revealed significant differences [p = 0.016], such that a greater percentage of ^4^He rats exhibited performance that was below the 5^th^ worst percentile performance of Sham rats than GCR rats. Lastly, the risk ratio calculator determined the absolute risk increase to be 37.00% (CI: 26.43% to 47.57%) for ^4^He rats with numbers needed to harm revealing that one out of every three (CI: 2.5 to 3.8) rats exposed to 10 cGy ^4^He will experience an increase in approach time. An absolute risk increase of 20% (CI: 10.50% to 29.50%) was observed for GCRsim rats with one out of every 5 rats exposed to 10 cGy GCRsim expected to exhibit an increased time to approach the string.

#### 3.1.2. Pull Duration

The duration it took rats to pull in the string to reach a piece of Cheerio tied at the end was evaluated before SR exposure, 72 hours after irradiation, and the change in performance between the two time points (see Figure 3). Prior to SR exposure, Sham (M = 32.378, SD = 52.571), 4He (M = 40.104, SD = 56.485), and GCRsim (M = 57.849, SD = 60.067) rats took similar amounts of time to pull in the string as revealed by an ANOVA [F (2, 57) = 0.924, p = 0.403, η^2^_p_ = 0.033]. Following irradiation, Mann Whitney tests also failed to reveal significant group differences between either ^4^He- [p = 0.077, M = 30.608, SD = 43.214] or GCRsim-exposed [p = 0.145, M = 28.236, SD = 49.835] rats relative to Sham rats (M = 9.976, SD = 4.5 06: see Figure 2E).

However, rats exhibited within group variability in pull duration. Thus, a KDE analysis of pull duration also calculated the 5^th^ worst percentile for Sham rats to be 20.07 s (see Figure 2F). The percentage of SR-exposed rats that had performance below the 5^th^ percentile of the Sham rats’ performance was 59% for ^4^He- and 50% for GCRsim exposed rats. Fisher’s exact test comparing pull duration for Sham and ^4^He-exposed rats revealed statistically significant differences between groups [p < 0.001]. In addition, pull duration for GCRsim rats significantly differed from Sham rats [Fisher’s exact: p < 0.001], while irradiated groups failed to differ from each other [p = 0.256]. Both ^4^He- and GCRsim-exposed groups had a significant percentage of rats that fell below the 5^th^ worst percentile of Sham rats, exhibiting longer pull durations to reach the Cheerio tied at the end of the string 72 hours after irradiation exposure. Subsequently, the KDE generated data that was entered into a risk ratio calculator determined both level of absolute risk increase and numbers needed to harm for ^4^He rats to be 54% (CI: 43.46% to 64.54%) and 2 (CI: 1.5 to 2.3) and GCRsim rats to be 45% (CI: 34.31% to 55.69%) and 2 (CI: 1.8 to 2.9), respectively. Thus, for every 2 rats exposed to 10 cGy of ^4^He or GCRsim, one is expected to take longer to pull in the string to reach the Cheerio tied at the end.

Next, the change in pull duration was compared between Groups (see Figure 3C). An ANOVA failed to reveal a significant effect of Group [F (2, 56) = 0.990, p = 0.378, η^2^_p_ = 0.035] between Sham (M = −22.401, SD = 52.739), ^4^He (M = −3.853, SD = 64.020), and GCR (M = −29.869, SD = 67.965) rats. KDE analyses calculated the 5^th^ worst percentile of Sham rats’ performance to be 28.8 seconds for the change in pull duration from between conditions, with 25% of ^4^He and 8% of GCR rats below this performance level. Fischer’s exact tests revealed that a significant proportion of ^4^He rats [p = 0.0001] fell below this 5^th^ percentile, while GCR rats did not significantly differ from Sham rats [p = 0.568]. However, irradiated groups significantly differed [p = 0.002] from each other with a greater percentage of ^4^He rats falling below the 5^th^ worst percentile performance of Sham rats than GCR rats.

### 3.2. Characterization of movement accuracy

Rats engaged in subcomponents (Lift, Advance, Grasp, Pull, Push) of movement to pull in the string (see Figure 4) with a mixture of contacts and misses. Following ^4^He-exposure, irradiated rats exhibited changes in movement accuracy ratio, or contacts and misses made with both the hands and mouth (see Figure 6).

**Figure 4:**
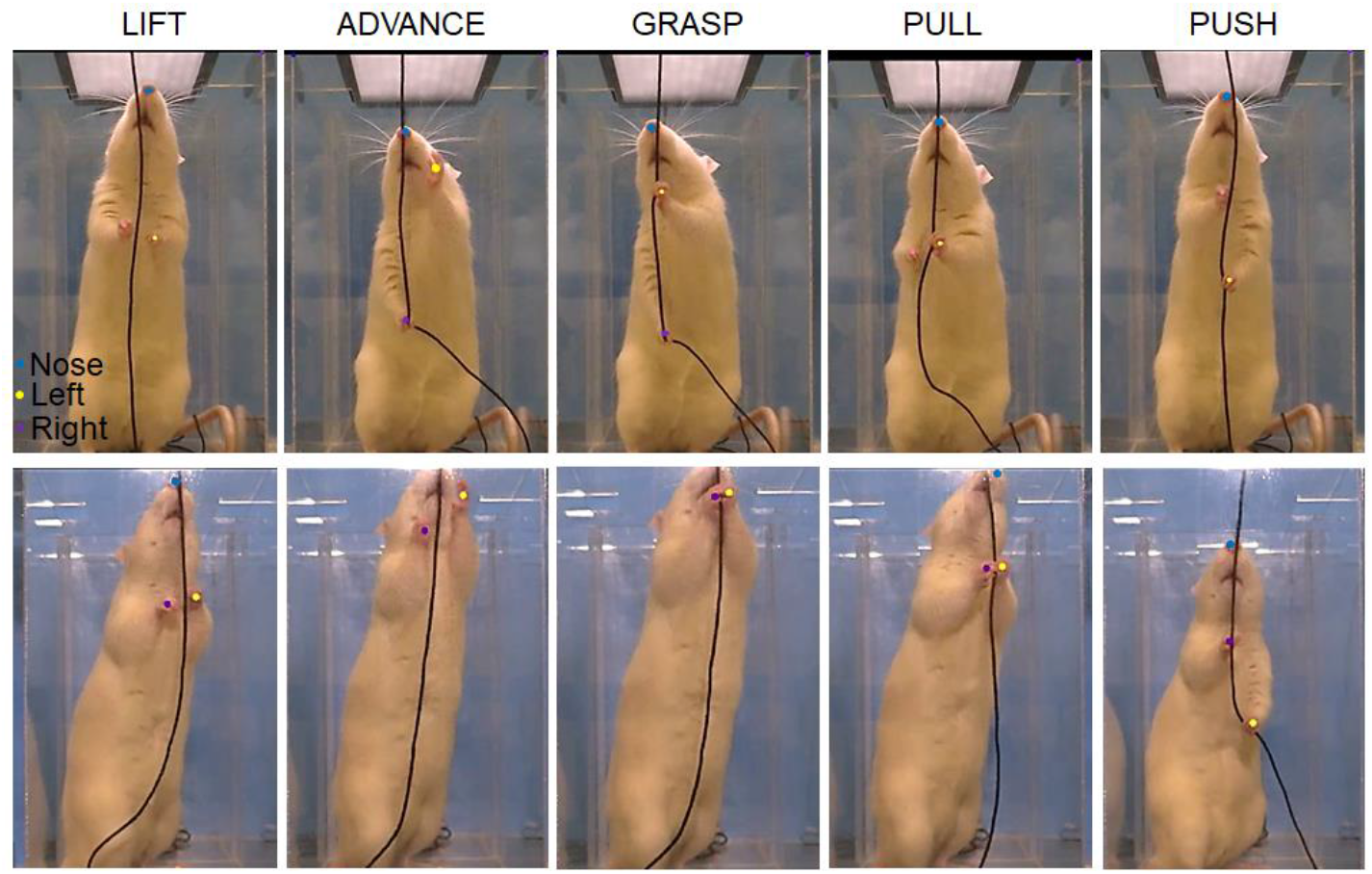
Subcomponents of string-pulling behavior are shown for a rat: Lift, Advance, Grasp, Pull, and Push. The top set of images demonstrate independent reach and withdraw movement where the hands alternate directions, while the bottom pictures show simultaneous reach and withdraw hand movement moving in the same direction. The rats body parts were labeled by DLC (V2.1 [22]; nose: blue, left: yellow, and right: purple).

**Figure 5:**
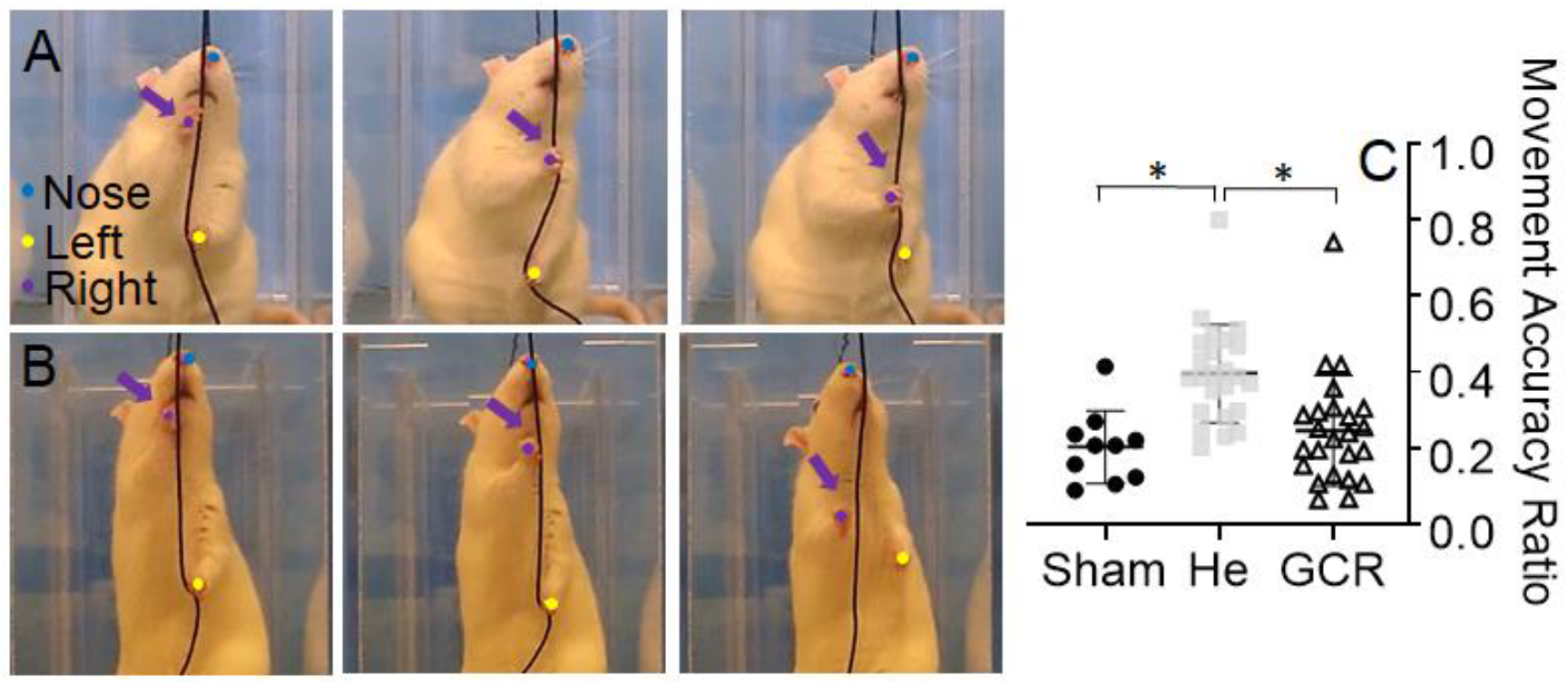
A frame-by-frame right-hand contact is shown for a rat (A) and a miss (B). The movement accuracy ratio conducted on misses and contacts (C) revealed that ^4^He-exposed rats exhibited more misses and less contacts relative to both Sham and GCRsim rats. The rats body parts were labeled by DLC (nose: blue, left: yellow, and right: purple). *p < 0.050

**Figure 6:**
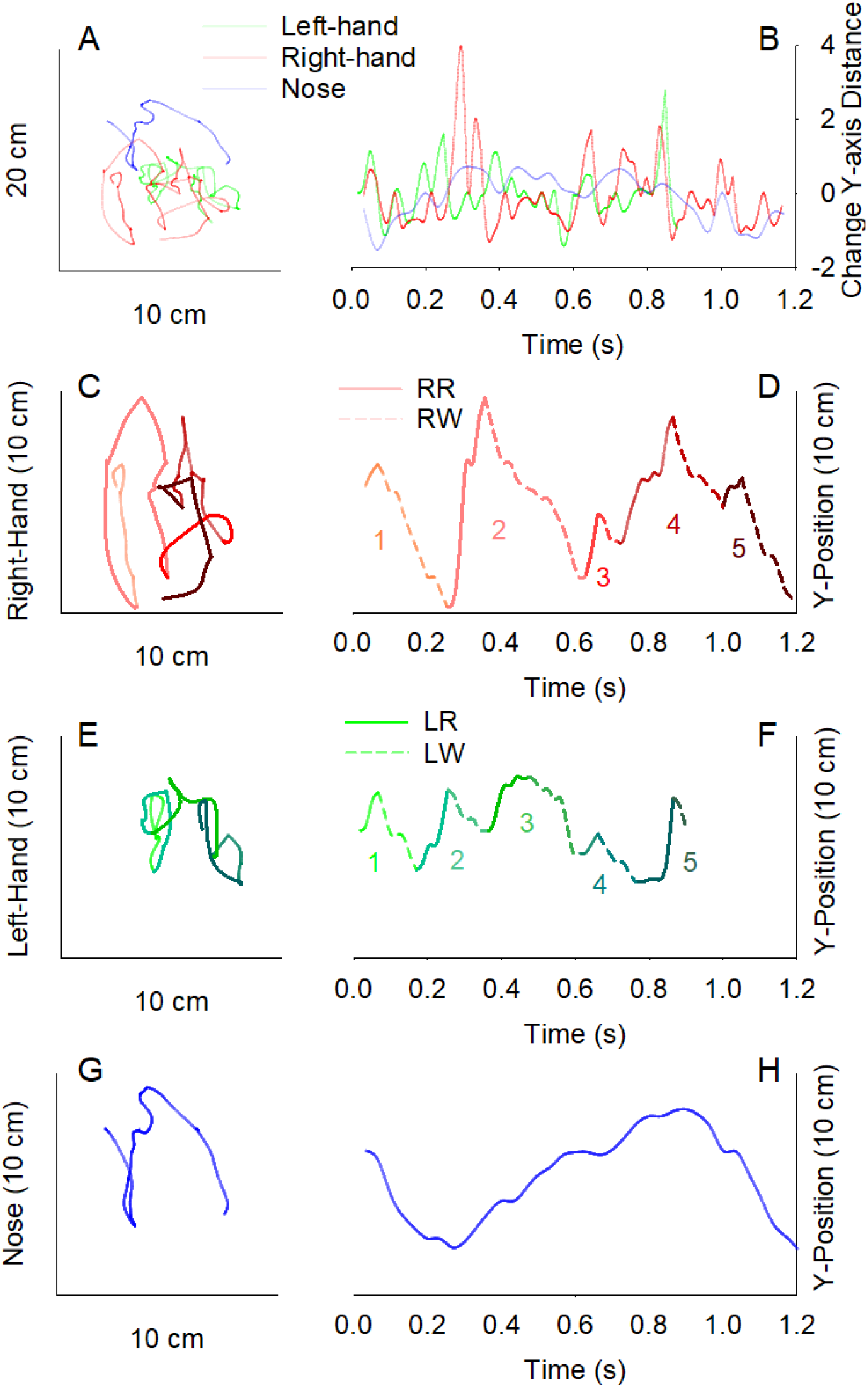
Topography of both hands and the nose are displayed for the first five cycles (one cycle is one reach and one withdraw) of pulling for a rat (A). The associated change in distance moved in the y-axis that was used to segment movement into upward reaches and downward withdraws is shown for a rat (B). Right-hand movement is color-coded by pulling cycle for the topography (C) and associated Y-position in space (D). A breakdown of the left-hand topography (E) and Y-position (F) and the nose (G-H) is also shown.

An ANOVA that was used to evaluate movement accuracy revealed a significant effect of Group [F (2, 54) = 8.145, p < 0.001, η^2^_p_ = 0.229]. Post hoc analysis (Tukey HSD < 0.050) demonstrated that ^4^He-exposed (M = 0.385, SD = 0.135) rats exhibited a greater proportion of misses and less contacts relative to both Sham (M = 0.219, SD = 0.106) and GCRsim-exposed (M = 0.249, SD = 0.151) rats. ^4^He rats experienced more issues contacting the string than Sham rats at this acute time point.

### 3.3. Kinematic analyses

String-pulling behavior consists of alternating upward reach and downward withdraw movements of the hands along with nose movement (see Figure 6). The organization of these movements is influenced by changes in hand and mouth contacts and misses, tracking of the string with the nose/vibrissae, and posture. Reach and withdraw components were characterized by trajectories that displayed varying distances and speeds traveled across bouts with peak speeds occurring at the initiation of rat string-pulling behavior (see Figure 7). Yet no group differences were observed in distance traveled or peak speed for reaches 72 hours after irradiation exposure (see Table 1).

**Table 1.**
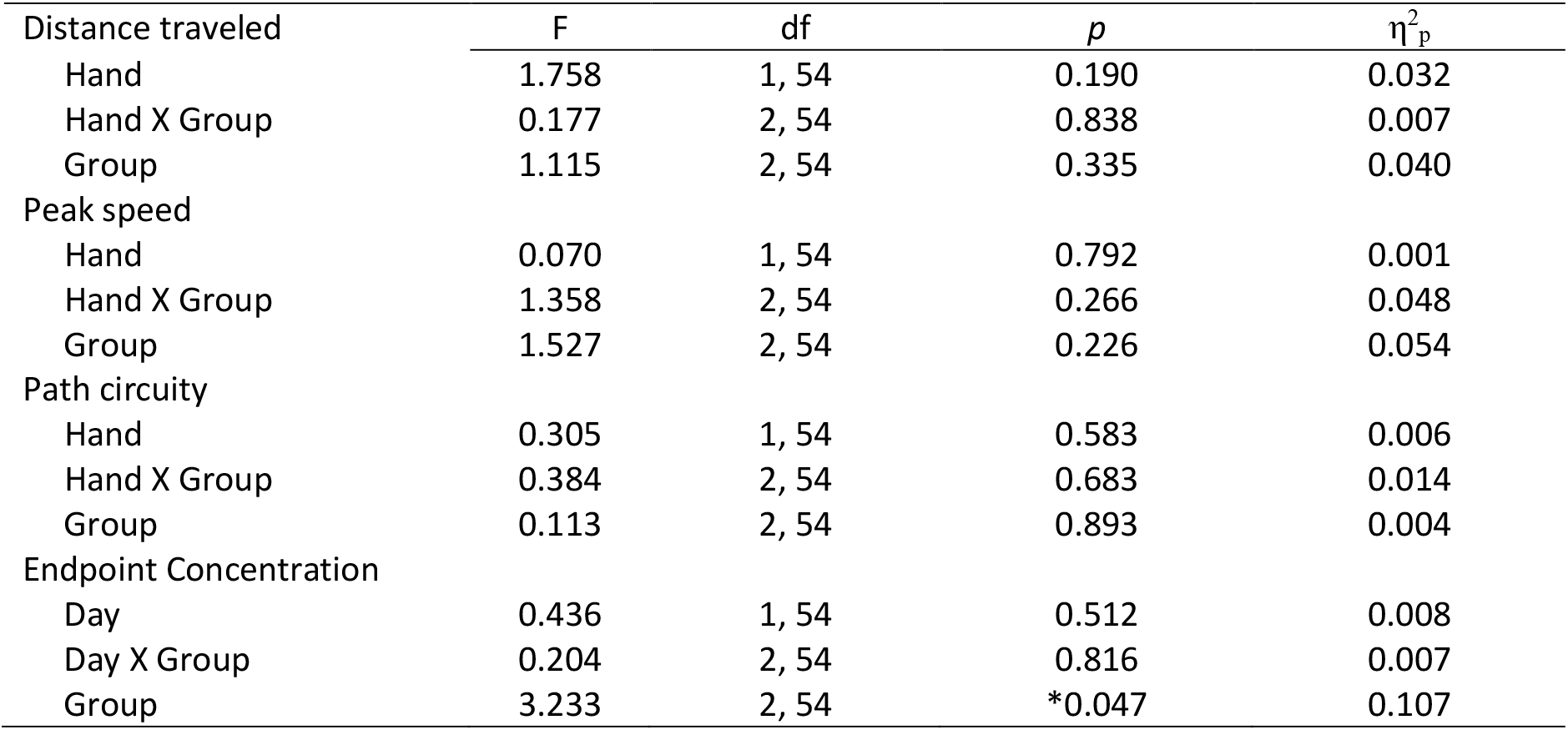
Left- and right-hand reach kinematics.

**Figure 7:**
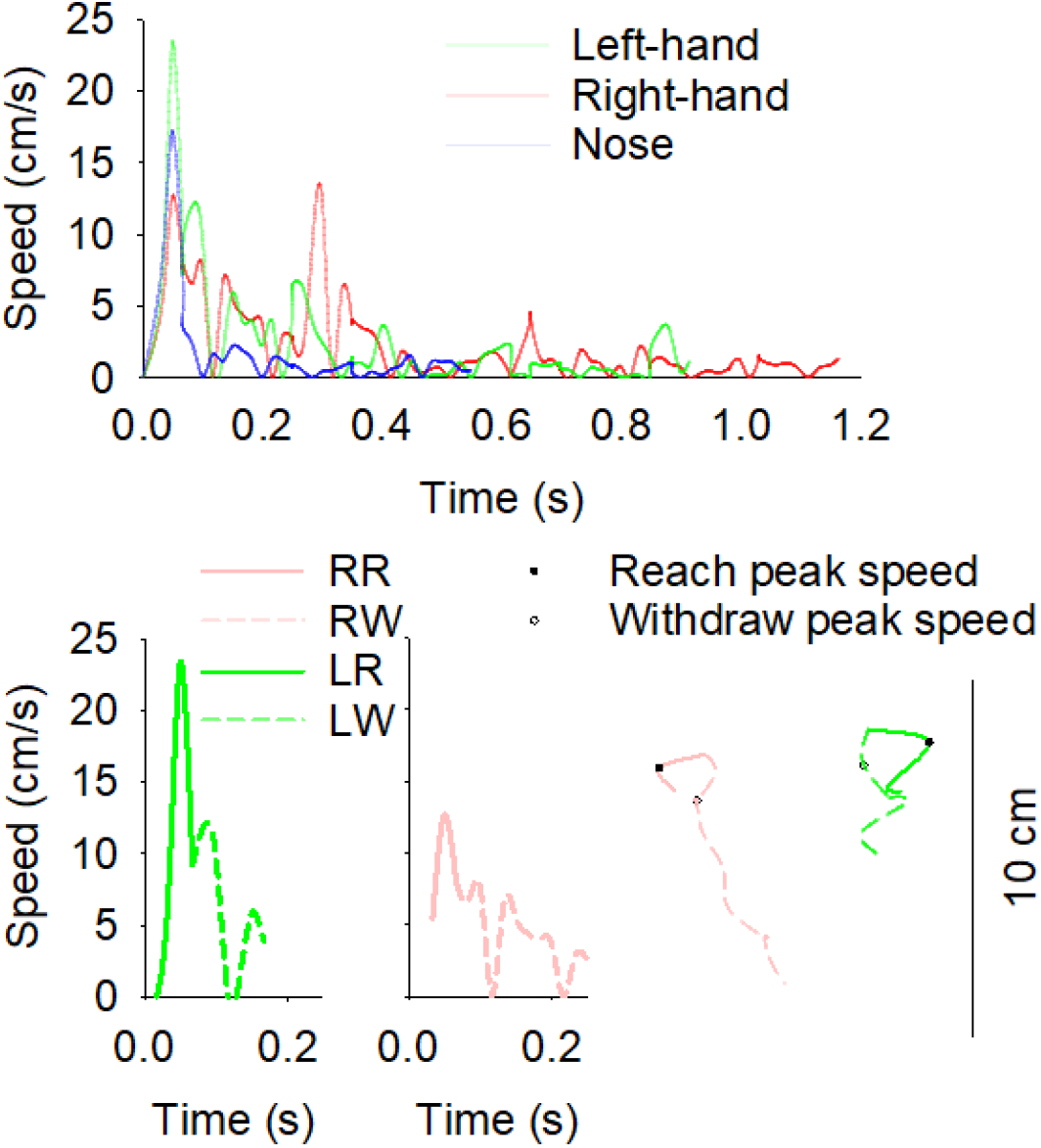
Representative moment-to-moment speeds are plotted for the hands and the mouth of the first five pulling cycles for a rat (top). The speeds are plotted for the first reach and withdraw in the bout of string-pulling (bottom left) with the associated topography (bottom right).

During reaches, rats lifted their hands up to aim, advance toward, and then grasp to contact or miss the string (see Figure 8). The kinematics and topography of these reaching movements were organized similarly between rats during string-pulling behavior with one exception (see Table 1). While reach path circuity was also similar between groups 72 hours after irradiation (see Figure 9), a repeated measures ANOVA conducted on the concentration of left- and right-hand endpoints exclusively revealed a significant effect of Group [F (2, 54) = 3.233, p = 0.047, η^2^_p_ = 0.107] (see Figure 10). Post hoc analysis revealed that ^4^He-exposed rats exhibited less concentrated reach endpoints relative to GCRsim rats (Tukey HSD < 0.050). Due to significance in reach endpoint concentration (first order), heading direction (second order) could not be statistically evaluated for reaches.

**Figure 8:**
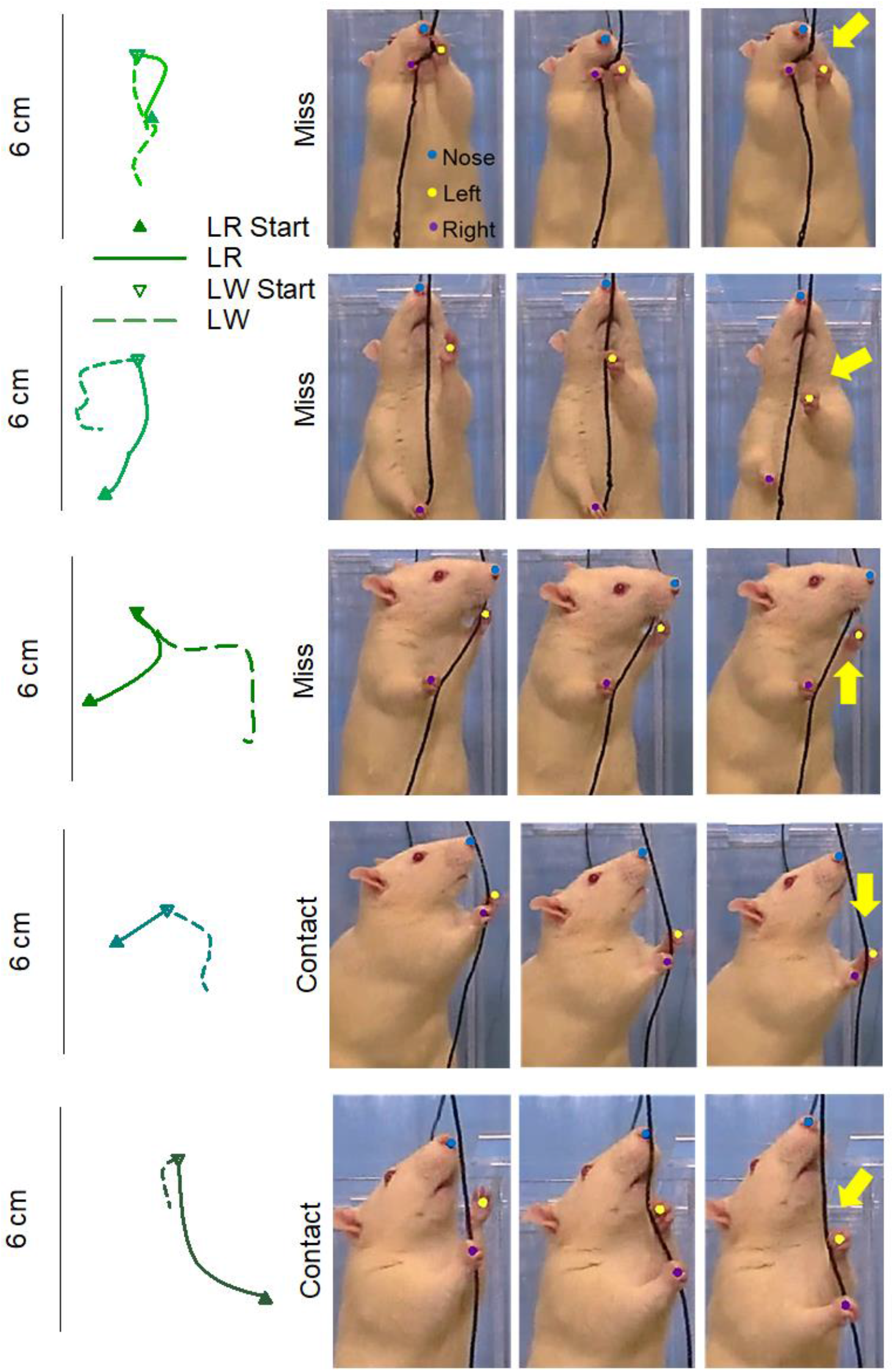
Reach and withdraw topography are displayed for the first five cycles of pulling for a rat on the left. The associated contacts or misses that occurred during each of these cycles is depicted on the right.

**Figure 9:**
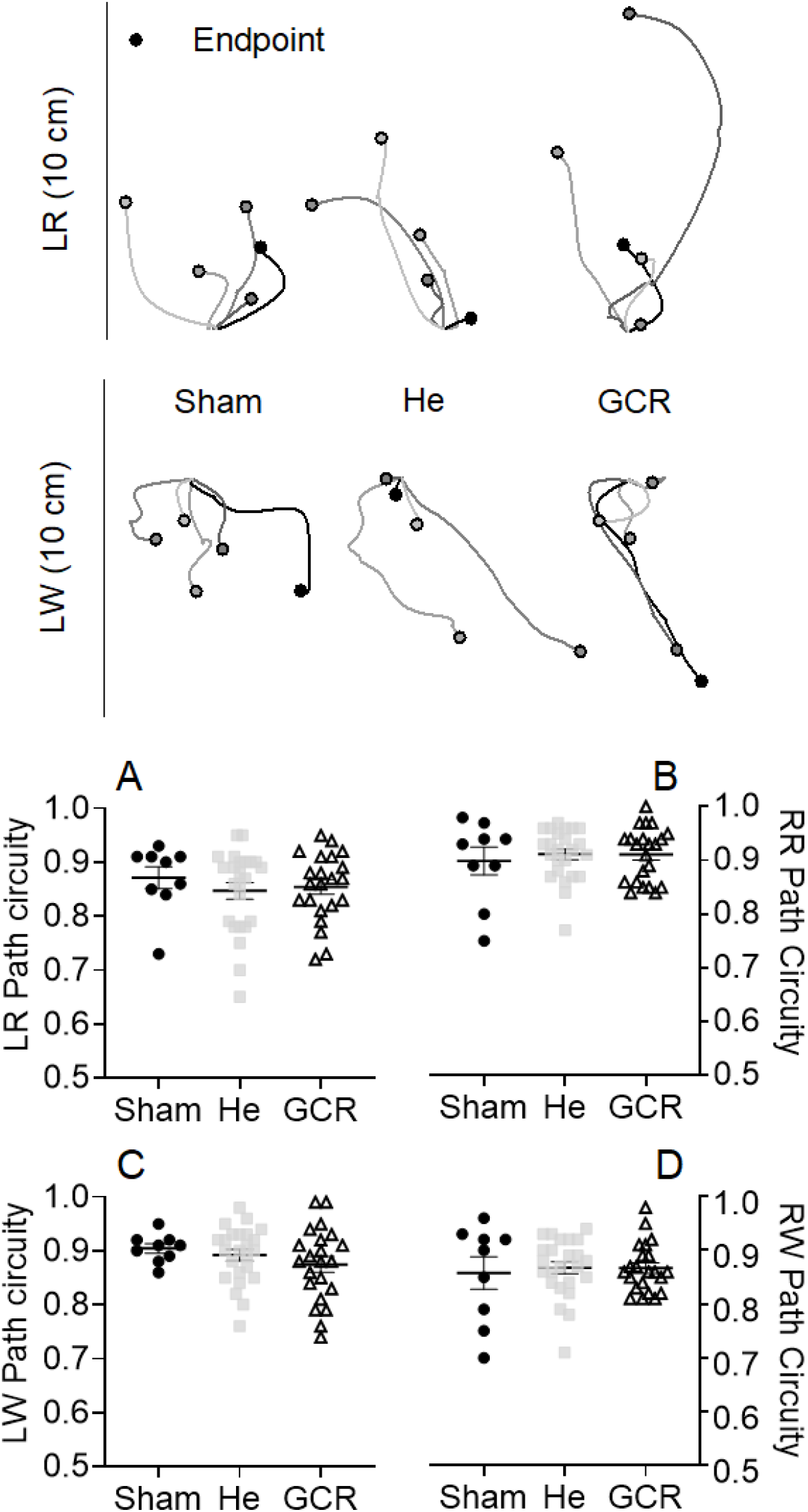
Transformed reach (left) and withdraw (right) trajectories are displayed for the left hand for a representative rat from each group. No significant differences were observed in reach path circuity with the left-(A) and right-hand (B) or withdraw path circuity for the left-(C) and right-hand (D).

**Figure 10:**
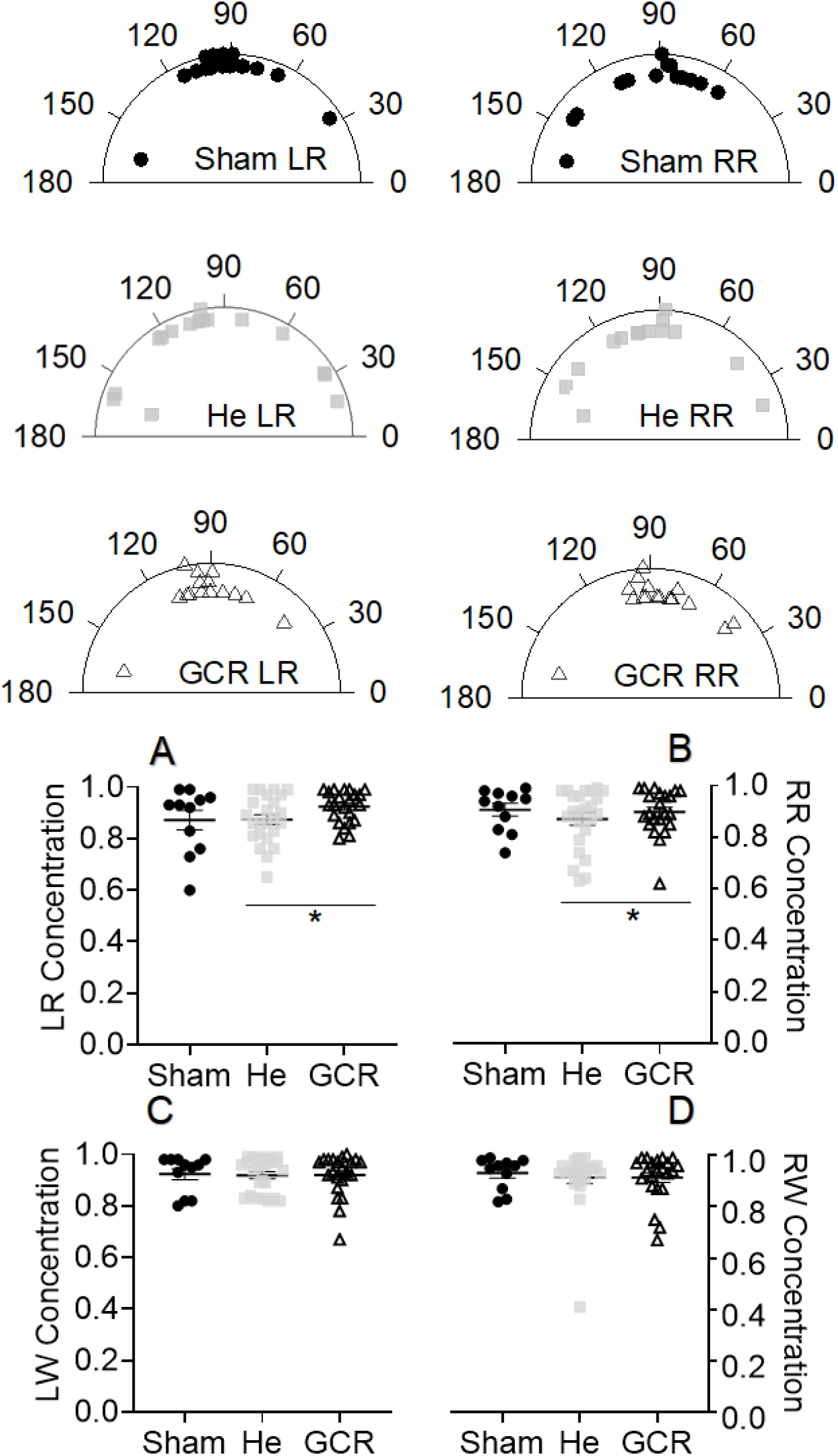
The concentration of reach endpoints is shown for a representative rat from each group. ^4^He-exposed rats exhibited less concentrated reach endpoints with both the left- (A) and right-hand (B) relative to rats exposed to GCRsim. No differences were observed in the concentration of left-or right-hand withdraw endpoints. *p < 0.050

After grasping the string, all rats lowered their arm and hand to pull then push the string into the apparatus. Unlike reaching movements, measures characterizing withdraw kinematics failed to reveal any significant differences between groups 72 hours after irradiation (see Table 2). Analyses of kinematic measures for the nose also revealed non-significant results for peak speed [F (2, 54) = 0.604, p = 0.550, η^2^_p_ = 0.022], and X [F (2, 54) = 0.014, p = 0.986, η^2^_p_ = 5.103e-4] and Y [F (2, 54) = 1.090, p = 0.343, η^2^ = 0.039] range of movement. Concentration of reach endpoints were selectively impaired in ^4^He-compared to GCRsim-exposed rats, while all other kinematic measures for withdraw subcomponents and the nose were organized similarly between groups

**Table 2.** Left- and right-hand withdraw kinematics.

| Distance traveled | F | df | $p$ | $\eta^2_p$ |
| --- | --- | --- | --- | --- |
| Hand | 1.412 | 1, 54 | 0.240 | 0.025 |
| Hand X Group | 0.264 | 2, 54 | 0.769 | 0.010 |
| Group | 1.836 | 2, 54 | 0.169 | 0.064 |
| Peak speed |  |  |  |  |
| Hand | 0.003 | 1, 54 | 0.959 | 4.852e-5 |
| Hand X Group | 0.148 | 2, 54 | 0.863 | 0.005 |
| Group | 0.271 | 2, 54 | 0.763 | 0.010 |
| Path circuitry |  |  |  |  |
| Hand | 2.844 | 1, 54 | 0.097 | 0.050 |
| Hand X Group | 0.551 | 2, 54 | 0.579 | 0.020 |
| Group | 0.199 | 2, 54 | 0.820 | 0.007 |
| Endpoint concentration |  |  |  |  |
| Day | 8.422e-6 | 1, 54 | 0.977 | 1.560e-5 |
| Day X Group | 0.134 | 2, 54 | 0.874 | 0.005 |
| Group | 0.662 | 2, 54 | 0.520 | 0.024 |
| Heading direction |  |  |  |  |
| Day | 3.883 | 1, 54 | 0.054 | 0.067 |
| Day X Group | 0.182 | 2, 54 | 0.834 | 0.007 |
| Group | 0.268 | 2, 54 | 0.766 | 0.010 |

## 4. Discussion

This study is the first assessment of fine motor control to reveal performance deficits in a rodent model 72 hours after SR exposure. Thus, sensorimotor deficits are both an acute and late [19] reaction following SR exposure. Rats exposed to 10 cGy ^4^He exhibited multiple deficits in the string-pulling task 72 hours after exposure that were characterized by: 1) increased approach time, 2) longer pull durations, 3) greater misses and less contacts, and 4) less concentrated reach endpoints with both hands. GCRsim exposure selectively increased rat pull duration at this acute time point. These disruptions in string-pulling behavior provide evidence for impairments in sensorimotor function and motivation.

Behavioral assessments are typically conducted 1 to 3 months post-irradiation exposure. While several studies have reported acute CNS effects following SR exposure, to our knowledge, only one other study has evaluated performance related to sensorimotor function at an acute time point [5]. However, the current work establishes impaired function and suggests that SR impacts performance within a time frame that translates to in-flight missions for astronauts. This demonstrates the importance of evaluating performance in multiple systems during mission relevant time periods.

Deficits in fine motor control were recently identified for the first-time following SR exposure. Rats exposed to 5 cGy ^28^Si exhibited longer pull durations and a reduced ability to contact the string ~7 months after SR exposure [19] similar to SR-exposed rats assessed 72 hours after irradiation. While multiple changes in movement kinematics related to distance and direction of reach and withdraw subcomponents were observed in ^28^Si-exposed rats at a protracted time point, the same level of impairments were not present in the current study 72 hours after SR exposure. However, when ^28^Si-exposed rats were assessed ~7 months post-irradiation, they were more than twice the age (~15 mo) of the rats used in the current study (~ 7 mo). Therefore, protracted effects of SR may interact with age to account for the severity in string-pull performance deficits previously reported after ^28^Si exposure. Regardless, ^4^He-exposed rats exhibited less concentrated reach endpoints with both hands compared to GCRsim rats suggestive of disruptions in the consistency of reach-to-grasp trajectories that likely contribute to misses made by the hands. This may also suggest that there are ion-specific, temporal gradients, and/or aging interactions for the effects of SR exposure on string-pulling behavior.

Different types of information that guide the organization of string-pulling behavior may have been impacted by SR at this acute time point. Rats must first detect and approach the string which is typically first done with the vibrissae and nose followed by the hands and mouth. Then, rats must be motivated to engage in pulling behavior to retrieve a piece of Cheerio tied at the end of a string. Previous work has shown that pulling strings with different reinforcement rates (1/8 vs 4/8 unrewarded strings) or strings of different lengths (1.5 m and 3 m) distinctly influences approach time and pull duration, respectively, during longer, unrewarded probe strings [21]. However, no changes were observed in misses/contacts or kinematic measures of performance upon varying reward rate (i.e., motivational component) or string length (i.e., temporal component). While these factors were not exclusively assessed in the current study, evidence for motivational and/or attention deficits have been reported following exposure to SR in various behavioral tasks, including psychomotor vigilance task [31], attentional set shifting [20, 32–35], and social withdrawal [27]. Therefore, this work taken with the current findings that ^4^He rats take longer to approach the string, and both SR groups engage in longer pull durations, it is possible that irradiated rats may have experienced attentional, temporal, or motivational impairments that influenced performance in the string-pull task 72 hours after exposure.

The string-pulling task involves sensorimotor function and fine motor control of the hands and fingers to grasp a string. While increased misses and decreased contacts were observed by ^4^He rats in the current study, grasping motions were still made with the hands even when the string was missed. Similar, yet more severe deficits were observed after focal devascularization of the forelimb region within sensorimotor cortex, with rats exhibiting further distances traveled, faster peak speeds, and reduced endpoint concentrations in addition to increased pull duration and misses/hooks [24]. Sensorimotor dependent tasks that involve fine motor control also rely on the integration of information from sources that originate outside of the sensorimotor cortex, including motor afferent copies and spatial hand position that involves distance and direction.

Proprioception is one somatosensation that involves awareness of the body in space and time [36–37] that is important for the organization of appropriate limb coordination during movement [38]. It is possible that ^4^He-exposed rats experienced disruptions in proprioception 72 hours post-irradiation, and this may provide an explanation for increased misses and changes in reach endpoint concentrations. Both peripheral and/or central nervous system damage from whole body SR exposure has the potential to impair proprioception, since this sensory information arises from the peripheral system and feeds into the central system [39]. Proprioceptive information in the brain comes from the posterior parietal cortex which generates information about hand movements, including distance and direction estimation. Therefore, damage to this cortical area or its connections has the potential to disrupt the heading direction of reach endpoints and contribute to misses. While proprioceptive deficits have been reported following long duration stays on the International Space Station where astronauts are exposed to microgravity [40–41], not much is currently known about the effects of SR alone on proprioception. If exposure to SR impacts proprioception, then additive disruptive effects may occur on deep space missions where astronauts will experience multiple space flight stressors, including SR exposure and microgravity. However, it is also likely that changes to multiple systems involved in the organization of string-pulling behavior influenced the deficits observed 72 hours after SR exposure. Therefore, behavioral assessments that evaluate multiple aspects of performance (i.e., cognition and sensorimotor function) that are important for mission success should be used to characterize the effects of SR and other space flight stressors.

In this study, rats were exposed to the same dose (10 cGy) of two types of SR: one consisting of ^4^He ions and the other of a composite of several ions and energies. Regardless of SR exposure type, impaired string-pulling performance was present 72 hours after irradiation. However, at this acute time point, exposure to ^4^He resulted in greater deficits than 6-ion GCRsim, including longer approach times, increased misses with less contacts, and reduced concentration of reach endpoints. While there are multiple possible explanations for these differential effects, at this time these are largely speculative due to the lack of data on the ion/LET dependency. Only eight published studies [4, 15–16, 42–46] currently exist on the impact of multi-ion GCR exposure that mostly assess cognitive function. Aside from the present work, one study has directly compared this multi-ion spectrum to a single ion in a touchscreen associative recognition memory task where rats exposed to GCRsim exhibited greater deficits when presented with changes in multiple stimuli at once relative to ^4^He rats [4]. Overall, the current published data suggests that depending on the behavioral end point investigated, the impact of the 6-ion GCRSim on cognitive performance was either the same or opposite to those induced by a Z<15 multi-ion beam [18]. While string-pulling deficits at this acute time point were more pronounced after exposure to ^4^He than 6-ion GCRsim, performance measures on tasks of cognitive and sensorimotor function may not be equally impacted. Thus, the comorbidity of sensorimotor and cognitive impairments needs to be examined. Further, how such performance deficits change over time is also not currently known. Therefore, it is likely that SR exposure has complex multidimensions influencing the pattern of observable deficits which may be task dependent.

## 5. Conclusions

The present study is the first to identify deficits in sensorimotor function and motivation, temporal, and/or attentional processes 72 hours after SR exposure. Overall, rats exposed to ^4^He exhibited more misses and less contacts relative to both Sham and GCRsim-exposed rats. ^4^He-exposed rats also demonstrated less concentrated reach endpoints with both hands relative to GCRsim rats which likely contributed to misses. Lastly, increased time to approach (^4^He) and pull (^4^He and GCR) the string following SR exposure further supports disruptions in motivation, temporal, and/or attentional processing. While exposure to GCRsim did not have the same impact as ^4^He at this time point, additional deficits may develop over time with different temporal trajectories.

## Acknowledgements

This work was funded by NASA grant support NNX14AE73G and NNX16AC40G. The authors are thankful to Dr. Adam Rusek, Paula Bennett, and Deborah Snyder, at NSRL and the BLAF staff for their help with the irradiation of the rats at Brookhaven National Laboratories and animal care. This study would not have been possible without their help.

